# BrainPalmSeq: A curated RNA-seq database of palmitoylating and de-palmitoylating enzyme expression in the mouse brain

**DOI:** 10.1101/2021.11.23.468539

**Authors:** AR Wild, PW Hogg, S Flibotte, G Nasseri, R Hollman, K Haas, SX Bamji

## Abstract

Protein *S*-palmitoylation is a reversible post-translational lipid modification that plays a critical role in neuronal development and plasticity, while dysregulated *S*-palmitoylation underlies a number of severe neurological disorders. Dynamic *S*-palmitoylation is regulated by a large family of ZDHHC palmitoylating enzymes, their accessory proteins, and a small number of known de-palmitoylating enzymes. Here, we curated and analyzed expression data for the proteins that mediate *S*-palmitoylation from publicly available RNAseq datasets, providing a comprehensive overview of their distribution in the mouse nervous system. We developed a web-tool that enables interactive visualization of the expression patterns for these proteins in the nervous system (http://brainpalmseq.med.ubc.ca/), and explored this resource to find region and cell-type specific expression patterns that give insight into the function of palmitoylating and de-palmitoylating enzymes in the brain and neurological disorders. We found coordinated expression of ZDHHC enzymes with their accessory proteins, de-palmitoylating enzymes and other brain-expressed genes that included an enrichment of *S*-palmitoylation substrates. Finally, we utilized ZDHHC expression patterns to predict and validate palmitoylating enzyme-substrate interactions.

## Introduction

Protein *S-*palmitoylation is a post-translational lipid modification that mediates dynamic changes in protein stability, function, and membrane localization. *S-*palmitoylation is defined as the reversible formation of a cysteine residue thioester bond with the fatty acid palmitate, and is the most prevalent post-translational lipid modification in the brain. Dynamic changes in *S*-palmitoylation are critical for neuronal development and synaptic plasticity (Fukata et al., 2013; Fukata and Fukata, 2010; Globa and Bamji, 2017; Matt et al., 2019), oligodendrocyte differentiation and myelination (Ma et al., 2021; Schneider et al., 2005), and astrocyte proliferation (Yuan et al., 2021). Furthermore, numerous neurological and psychiatric diseases have now been attributed to mutations in the genes encoding palmitoylating and de-palmitoylating enzymes, including schizophrenia, intellectual disability and CLN1 disease (Mukai et al., 2004; Nita et al., 2016; Raymond et al., 2007), underscoring the importance of proper regulation of *S*-palmitoylation for normal brain function.

*S*-Palmitoylation is mediated by a family of ZDHHC enzymes that share a consensus ‘Asp-His-His-Cys’ catalytic domain. These enzymes are structurally heterogeneous multi-pass transmembrane proteins that localize to a variety of intracellular compartments, including the Golgi apparatus, endoplasmic reticulum (ER), recycling endosomes and the plasma membrane (Globa and Bamji, 2017). The ZDHHC enzymes are known to associate with accessory proteins that regulate their stability, activity, and trafficking (Salaun et al., 2020). Several de-palmitoylating enzymes have also been identified that act as the ‘erasers’ of *S*-palmitoylation, and are divided into three classes: the acyl-protein thioesterases that shuttle between the Golgi and cytosol (APTs; Vartak et al., 2014), the predominantly lysosomal palmitoyl-protein thioesterases (PPTs; Koster and Yoshii, 2019) and the more recently discovered α/β hydrolase domain-containing 17 proteins (ABHD17; Lin and Conibear, 2015). Unlike other post-translational modifications, palmitoylation lacks a consensus substrate amino sequence, and the mechanisms that govern ZDHHC enzyme-substrate interactions are controversial, with contrasting reports of substrate interactions being both promiscuous and specific (Malgapo and Linder, 2021). Currently these interactions are thought to be governed by the subcellular targeting of ZDHHCs enzymes and the presence of protein-protein interacting motifs within the ZDHHC N-and C-termini, which are highly diverse among the ZDHHC enzymes (Rana et al., 2018). Differential gene expression can also have a profound influence on protein interactions and may play a role in the coordination of *S*-palmitoylation in the brain. However, a detailed overview and analysis of the precise cellular and regional expression patterns of the palmitoylating and de-palmitoylating enzymes has not yet been described, and as such, little is known about how this expression is coordinated in the nervous system.

Recent advances in single-cell RNA sequencing (scRNAseq) techniques have enabled the classification of neuronal and non-neuronal cell types in unprecedented detail, providing a better understanding of cellular diversity and function in the nervous system, while also providing a means to study the expression patterns of individual genes across an ever-expanding range of brain regions and cellular classifications. Here, we capitalized on the recent surge in RNAseq publications characterizing regional and cellular transcriptomics of the mouse nervous system. We curated and analyzed expression data from a number of publicly available RNAseq mouse datasets to generate a detailed analysis of the expression patterns of the genes associated with *S*-palmitoylation in the mouse brain. Furthermore, we present an interactive web tool that allows user-driven interrogation of the expression patterns of palmitoylating and de-palmitoylating enzymes from numerous collated studies across a variety of brain regions and cell types (http://brainpalmseq.med.ubc.ca/). We demonstrate the utility of this resource by detailing the considerable cell type and regional heterogeneity in expression patterns of these enzyme families and their accessory proteins, revealing numerous cell type enrichments and co-expression patterns that allowed us to generate and test hypotheses about palmitoylating enzyme-substrate interactions.

## Results

### BrainPalmSeq: an interactive database to search palmitoylating and depalmitoylating enzyme expression in the mouse brain

The recent development of scRNAseq has revolutionized our understanding of the complex transcriptional diversity of neuronal and non-neuronal cell types in the brain. We found however, there were several barriers to the easy access for much of this data, with no single resource available to evaluate multi-study expression data. Data can also be difficult to access when studies are not accompanied by an interactive online web viewer, while datasets that do have a web viewer employ diverse interfaces that are often complex, particularly for large scRNAseq datasets. Furthermore, the differing study specific analysis pipelines, as well as the variety of data presentation formats in web viewers including heatmaps, bar charts, tables or t-SNE plots can make datasets difficult for non-bioinformaticians to interpret and compare. In order to remove these barriers and provide easy access to expression data for the proteins that regulate *S*-palmitoylation in the brain, we created ‘BrainPalmSeq’, an easy-to-use web platform allowing user driven interrogation of compiled multi-study expression data at cellular resolution through simple interactive heatmaps that are populated according to user selected brain regions, cell-types or genes of interest (http://brainpalmseq.med.ubc.ca/).

To create BrainPalmSeq we first curated three large datasets from whole-brain scRNAseq studies that were acquired through selection-free cell sampling to provide high resolution expression data covering hundreds of cell types at a variety of developmental ages (Rosenberg et al., 2018; Saunders et al., 2018; Zeisel et al., 2018). As scRNAseq has several caveats including low sensitivity and high frequency of dropout events leading to incomplete detection of expressed genes (Haque et al., 2017), we complemented these datasets where possible with curation of several bulk and pooled-cell RNAseq studies that used population-level ensemble measurements from whole-brain and region-specific studies. We further included selected studies for the major glial cell types and data from the most comprehensive neuron specific study performed to date by the Allen Institute (Supplementary File 1). Together, the datasets curated in BrainPalmSeq cover all major regions of the mouse nervous system across a variety of regional and cellular resolutions.

Expression data were extracted from selected studies for the 24 mouse ZDHHC genes (*Zdhhc1*-*Zdhhc25*, while *Zdhhc10* is omitted), as well as the best characterized de-palmitoylating enzymes (*Ppt1*, *Lypla1*, *Lypla2*, *Abhd17a*, *Abhd17b* and *Abhd17c*) and ZDHHC accessory proteins (*Golga7*, *Golga7b* and *Selk*). Where possible, data were processed from the raw transcripts or unique molecular identifier (UMI) counts using the same normalization protocol to allow for more consistent evaluation of differences in gene expression within datasets. We sampled from RNAseq datasets that used a diverse range of sample collection, processing and analysis techniques, allowing for direct visualization of the relative expression patterns of selected genes can be directly visualized within datasets. Users can then validate their observations across complimentary whole brain or region/cell-type specific datasets included in BrainPalmSeq. Dropdown menus allow for selection of individual ZDHHC genes or brain regions within each dataset, while the hover tool reveals metadata for each cell type, including neurotransmitter designations and marker genes. We provide download links to all expression data including cell type metadata so that users can replot gene expression profiles in their preferred format. To demonstrate the utility of this resource, we performed a detailed exploration of selected datasets from BrainPalmSeq, revealing how expression patterns can give insights into the function of the palmitoylating and de-palmitoylating enzymes in the mouse brain.

### ZDHHC expression in the nervous system shows regional and cell-type specific patterning

We began by exploring BrainPalmSeq data curated from the ‘MouseBrain’ dataset, which provides the broadest overview of expression patterns in the nervous system (Zeisel et al., 2018). This scRNAseq study sampled multiple dissected regions from the adolescent (mean age ∼P25) mouse central and peripheral nervous systems (CNS and PNS, respectively), identifying 265 transcriptomically unique cell-types (referred to herein as metacells) for which we plotted re-normalized ZDHHC expression values, according to the hierarchical cell-type clustering established by the original study (Figure 1A). While ZDHHC expression was detected in all regions of the nervous system, expression of the 24 ZDHHC genes was highly variable across metacell types and clusters. We measured the mean ZDHHC expression within each cluster to gain insight into which cell-types in the nervous system have the greatest overall expression of palmitoylating enzymes. The heatmap rows and columns were ranked (sorted by descending averages) to determine which cell-types had the highest expression of ZDHHCs, and which ZDHHCs were most abundantly expressed across cell-types (Figure 1B). Mean ZDHHC expression was particularly high in neurons of the PNS, along with cholinergic/monoaminergic and hindbrain neurons of the CNS. Of the non-neuronal metacell clusters, oligodendrocytes had the highest ZDHHC expression, while other glial cell-types appear at the lower end of the ranking (Figure 1B). *Zdhhc20* was the most abundantly expressed ZDHHC, with the highest mean expression across all cell-type clusters, followed by *Zdhhc2*, *Zdhhc17*, *Zdhhc3* and *Zdhhc21*, while expression of *Zdhhc11*, *Zdhhc19* and *Zdhhc25* were negligible. We next clustered neuronal metacells of the PNS and CNS according to the neurotransmitter expression combinations, revealing the highest mean ZDHHC expression was observed in neurons that utilized acetylcholine and nitric oxide as co-neurotransmitters, with cholinergic neurons featuring near the top of the list in several neurotransmitter combinations (Figure 1C). Monoaminergic neurons utilizing noradrenaline and serotonin also generally ranked high in the list, consistent with the data in Figure 1B that ranked cholinergic and monoaminergic neurons as the metacell cluster with the highest CNS ZDHHC expression overall, indicating a higher propensity for these cell-types to utilize *S*-palmitoylation as a post-translational mechanism to modify cellular signaling. We performed comparative analysis of ZDHHC expression on another large-scale scRNAseq study of the mouse brain that sampled a variety of cortical and subcortical structures of the adult mouse (P60-P70) (Saunders et al., 2018; ‘DropViz’; Figure 1-figure supplement 1A). We found expression patterns and enrichments to be similar across these two independent, large scale scRNAseq studies, supporting the general trends observed within the MouseBrain dataset.

**Figure 1.**
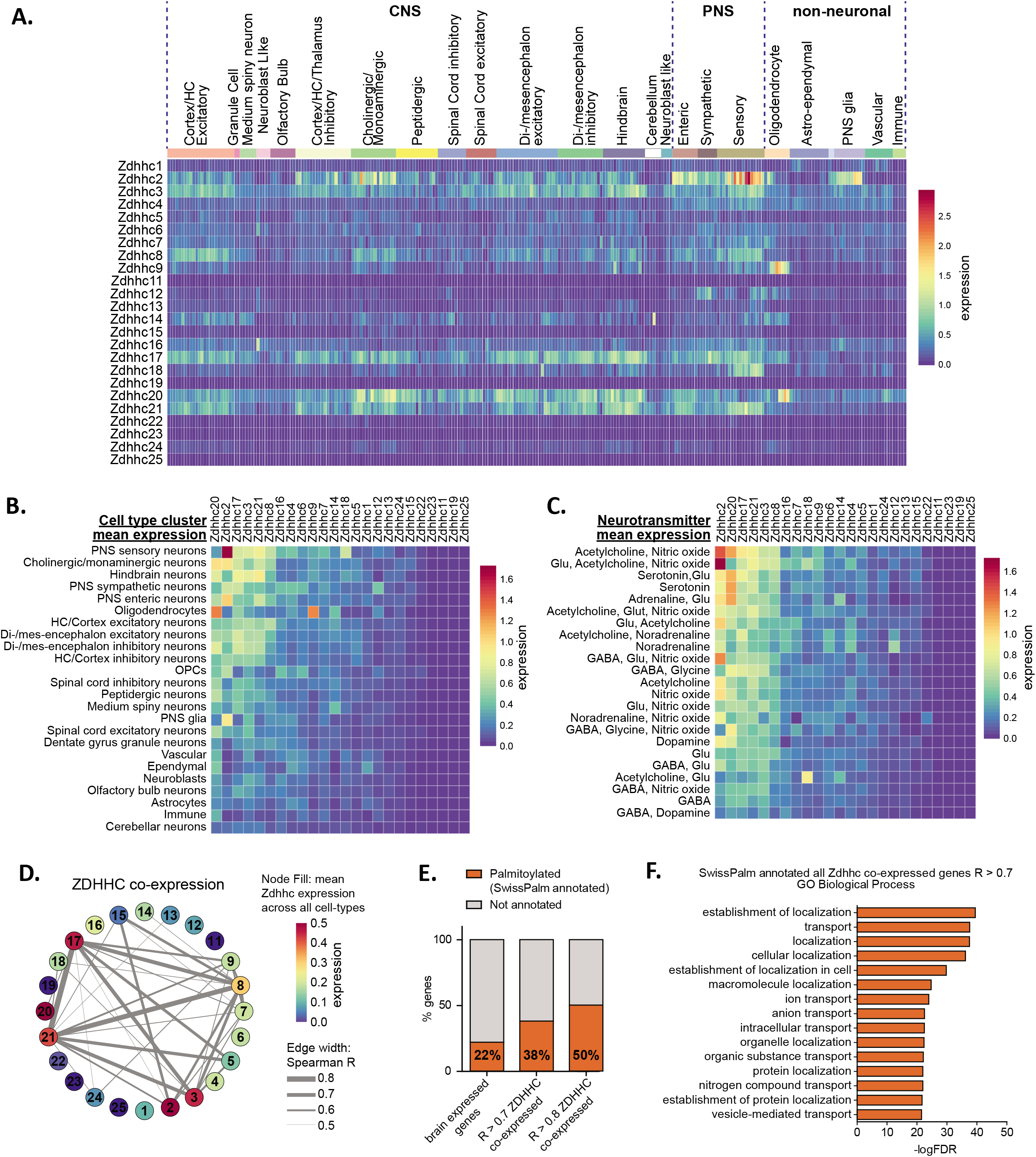
Heterogeneous ZDHHC expression in the mouse nervous system. (A) Heatmap showing expression for the 24 ZDHHC genes, extracted from scRNAseq study of mouse CNS and PNS (Zeisel et al., 2018). Each column represents one of the 265 metacells classified in the study. Metacells are organized along x-axis according to hierarchical clustering designations generated by Zeisel et al. Full metadata for this study available on BrainPalmSeq. (B) Heatmap showing mean ZDHHC expression per hierarchical cluster, with columns and rows sorted by descending mean ZDHHC expression per row/column. (C) Heatmap showing mean ZDHHC expression per neurotransmitter cluster for all PNS and CNS neurons. Columns and rows are sorted as in B. (D) Correlation network showing ZDHHC co-expression across all metacells in ‘MouseBrain’ (Spearman R > 0.5). Numbers in nodes correspond to ZDHHC number. Node color represents mean expression across all metacells. Edge thickness represents strength of correlation. (E) Graph showing proportion of genes from ‘MouseBrain’ dataset that are co-expressed with one or more ZDHHC and also substrates for *S*-palmitoylation (SwissPalm annotated). ‘Brain expressed’ *n* = 15,389 protein coding genes expressed in the postnatal mouse brain, curated from the MGI RNAseq studies database. ‘R > 0.7 ZDHHC co-expressed’ *n* = 914 genes co-expressed with one or more ZDHHC (Spearman R > 0.7). ‘R > 0.8 ZDHHC co-expressed’ *n* = 167 genes co-expressed with one or more ZDHHC (Spearman R > 0.8). Brain expressed vs. R > 0.7: p < 0.001; R > 0.7 vs R > 0.8: p < 0.01; Fisher’s exact test. (F) Graph of GO biological process analysis. Gene IDs from the ‘MouseBrain’ dataset (Zeisel et al., 2018) that showed correlated expression with one or more ZDHHC (R > 0.7) and were also Uniprot reviewed and SwissPalm annotated were used as input. Units for all heatmaps in figure: mean log2(counts per 10,000 + 1).

To gain insight into the potential networks of ZDHHC enzymes that might work together to coordinate *S*-palmitoylation in different cell types we performed co-expression analysis (Spearman correlation) between ZDHHC genes across all 265 metacell types in the MouseBrain dataset (Figure 1D). Neuron enriched *Zdhhc3*, *Zdhhc8*, *Zdhhc17* and *Zdhhc21* formed the strongest network of co-expression associations, while glial cell enriched *Zdhhc2, Zdhhc9* and *Zdhhc20* formed less robust correlations with other ZDHHCs. Weaker correlations were observed across the 565 cell-types in the DropViz dataset, which may reflect the absence of the PNS neurons and glia in this study (Figure 1-figure supplement 1D).

In order to create a list of potential substrates for the ZDHHCs in the mouse nervous system, we expanded our co-expression analysis to include all expressed genes from the ‘MouseBrain’ dataset that had significant correlation (R > 0.7) with one or more ZDHHC. We identified 914 genes with expression patterns that were significantly correlated with ZDHHCs. This list was cross-referenced with the mouse SwissPalm database of *S*-palmitoylated substrates identified in at least one palmitoyl-proteome or experimentally validated (SwissPalm annotated; Blanc et al., 2015, 2019). We found that genes that showed correlated expression with a ZDHHC were significantly enriched with *S*-palmitoylation substrates, indicating that ZDHHCs are more likely to be co-expressed with their *S*-palmitoylation substrates in the brain (Figure 1E). Co-expression analysis of the ‘DropViz’ dataset revealed a similar enrichment of *S*-palmitoylation substrates that were co-expressed with ZDHHCs (Figure 1-figure supplement 1E), supporting the notion of ZDHHC enzyme-substrate co-expression. PANTHER GO analysis of the ZDHHC co-expressed *S*-palmitoylation substrates curated from ‘MouseBrain’ revealed several significant enrichments in GO terms for biological processes related to protein localization (Figure 1F). These findings are consistent with the known role of *S*-palmitoylation in regulating protein localization and signaling complexes at cellular membranes.

### Heterogeneity in ZDHHC expression within excitatory neurons of the dorsal hippocampus

The hippocampus is a heavily studied brain region that is critical for learning and memory (Bird and Burgess, 2008). A recent pooled-cell RNAseq study of excitatory neurons in the hippocampus revealed extensive regional variability in gene expression profiles of the hippocampal tri-synaptic loop (hipposeq.janelia.org; Cembrowski et al., 2016). In order to clearly visualize if ZDHHC expression also varied within these different cell populations, we projected log transformed expression heatmaps generated in BrainPalmSeq for the ‘hipposeq’ dorsal-ventral excitatory neuron dataset on to anatomical maps of the dorsal hippocampus (Figure 2A). We observed considerable heterogeneity in the regional expression patterns of the ZDHHCs in the hippocampus. Hierarchical clustering analysis revealed that the ZDHHCs could be grouped into those that showed similar expression in all regions, those that were dentate gyrus granule cell (DG) enriched, DG depleted or CA1/CA2 enriched (Figure 2B). We generated comparative heatmaps for several scRNAseq studies curated in BrainPalmSeq that also quantified the hippocampal excitatory neuron transcriptome and found similar cross-study expression patterns for many of the ZDHHCs (Figure 2-figure supplement 1A). Furthermore, *in situ* hybridization data from the Allen Institute showed a high degree of overlap with the ‘hipposeq’ derived ZDHHC expression patterns, supporting the replicability of the expression patterns observed in the ‘hipposeq’ dataset (Figure 2-figure supplement 1B; Supplementary File 2).

**Figure 2.**
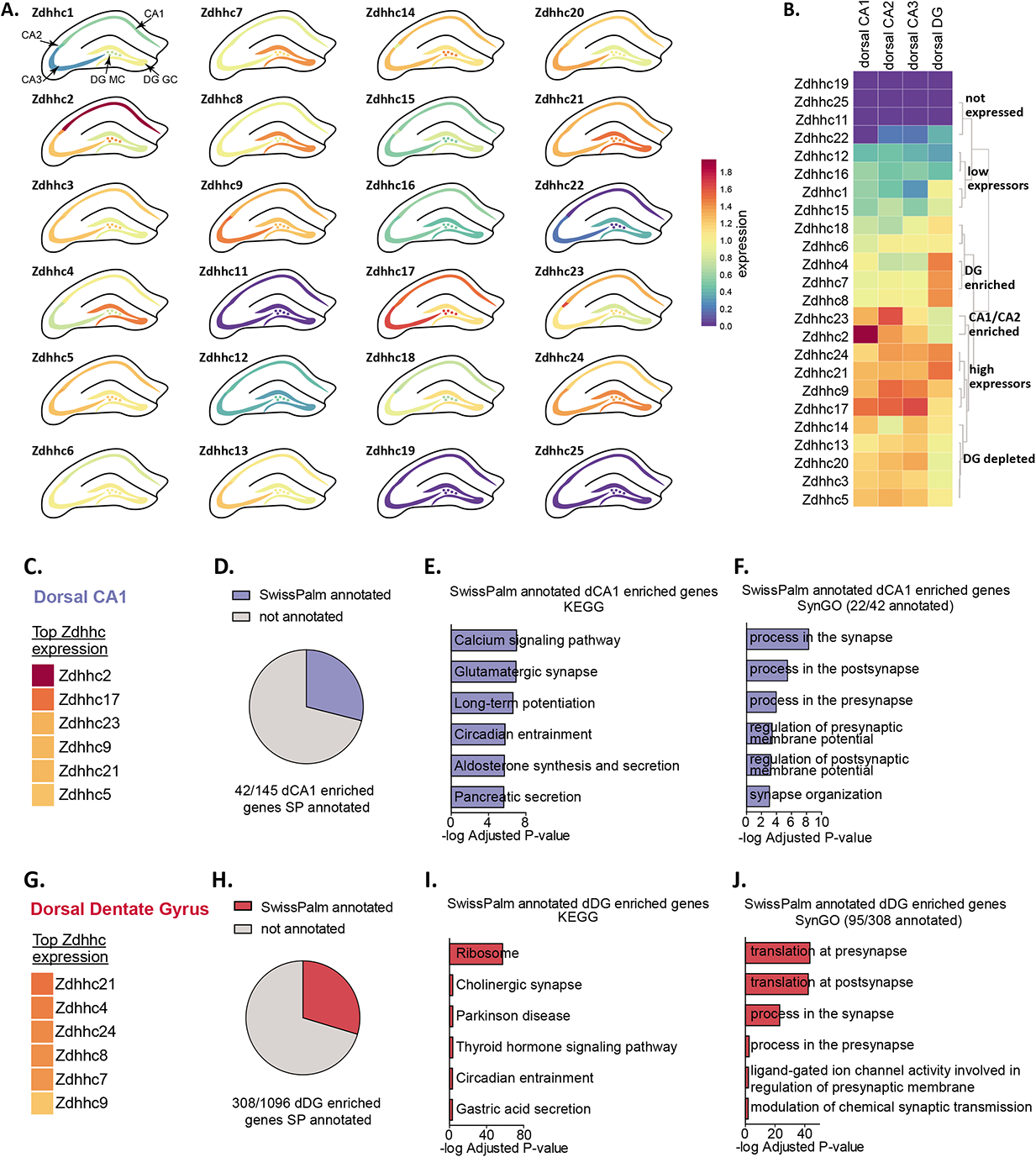
Diversity in ZDHHC expression and S-palmitoylation substrate expression in the hippocampus. (A) Heatmap of excitatory neuron ZDHHC expression from dorsal hippocampus (original pooled cell RNAseq data from Cembrowski et al., 2016) projected onto diagrams of dorsal hippocampus. (B) Hierarchical clustering of ZDHHC expression data in A. (C) Heatmap showing top 6 ranked expressing ZDHHCs in dorsal CA1 in descending order. (D) Pie chart showing proportion genes with enriched expression in dorsal CA1 (dCA1) that are also substrates for palmitoylation (SwissPalm annotated). (E) KEGG analysis of the dCA1 enriched/SwissPalm annotated genes. (F) SynGO analysis of the dCA1 enriched/SwissPalm annotated genes. (G-J) As in (C)-(F) but for the dorsal dentate gyrus (dDG). Heatmap legend in (A) applies to all heatmaps (logFKPM+1).

We next sought to utilize the ‘hipposeq’ dataset to determine if there might be regional differences in the expression of *S*-palmitoylation substrates in excitatory neurons of the dorsal hippocampus, which may be potential substrates for regionally enriched ZDHHC enzymes. To investigate the regionally enriched predicted hippocampal palmitoylome, we utilized the enrichment analysis tools built in to hipposeq.janelia.org (see Materials and Methods). Neurons in each hippocampal sub-region expressed unique *S*-palmitoylation substrates that were related to highly divergent functions. We found for CA1 neurons, which have the highest expression of *Zdhhc2*, *Zdhhc17*, *Zdhhc23* and *Zdhhc9* (Figure 2C), the CA1 enriched predicted palmitoylome (Figure 2D) generated KEGG pathways related to ‘Calcium signaling’, ‘Glutamatergic synapse’ and ‘Long term potentiation’, supporting the known role for *S*-palmitoylation in CA1 hippocampal synaptic plasticity (Figure 2E; Ji and Skup, 2021; Matt et al., 2019). The CA1 predicted palmitoylome was composed of around 46 % synaptic proteins (SynGO annotated), with SynGO ontologies related to ‘synaptic vesicle exocytosis’ and ‘synapse organization’ (Figure 2F; Koopmans et al., 2019). In contrast, the predicted palmitoylome of DG granule cells which have the highest expression of *Zdhhc21*, *Zdhhc4*, *Zdhhc24* and *Zdhhc8* (Figure 2G, H) generated KEGG pathways related to ‘Ribosome’, ‘Cholinergic synapse’ and ‘Parkinson’s disease’ (Figure 2I). The DG predicted-palmitoylome was composed of 29 % synaptic proteins (SynGO annotated), with SynGO ontologies related to ‘protein translation at presynapse’ and ‘protein translation at postsynapse’, revealing a potential role for palmitoylating enzymes in regulating translation in this cell-type that has not yet been studied (Figure 2J). Together, we have described patterns of restricted expression of ZDHHC enzymes and *S*-palmitoylation substrates in the dorsal mouse hippocampus, and generated regionally-enriched predicted-palmitoylomes that provide insight into the role of *S*-palmitoylation in neuronal function in each of these hippocampal sub-regions.

### Neocortical ZDHHC expression is partially segregated across cortical layers and neuronal subclasses

We next examined scRNAseq datasets curated in BrainPalmSeq from the cortex, beginning with a study of the primary somatosensory cortex (SSp; Zeisel et al., 2015). We projected heatmaps of pyramidal excitatory neuron ZDHHC expression generated in BrainPalmSeq onto cortical layer diagrams of SSp, again revealing anatomically heterogeneous excitatory neuron expression patterns for several of the ZDHHC transcripts. Clustering primarily grouped the enzymes according to expression levels, with *Zdhhc21*, *Zdhhc17* and *Zdhhc8* displaying the highest relative expression (Figure 3B). *Zdhhc2*, *Zdhhc3* and *Zdhhc20* expression was also high, with the remainder of the ZDHHCs having moderate to low expression. We compared these expression patterns with other datasets curated in BrainPalmSeq (Figure 3-figure supplement 1A), which revealed many consistent patterns of expression maintained across several independent studies. For example, multiple studies reported high expression of *Zdhhc8* in cortical Layer 2/3, enrichment of *Zdhhc2* in Layer 4 and elevated expression of *Zdhhc21* in all layers, particularly in Layer 5. *Zdhhc3* and *Zdhhc20* were also broadly expressed in all cortical layers across studies. Similar patterns were seen in the SSp region from available in situ hybridization studies from Allen Brain Institute (Figure 3-figure supplement 1B).

**Figure 3.**
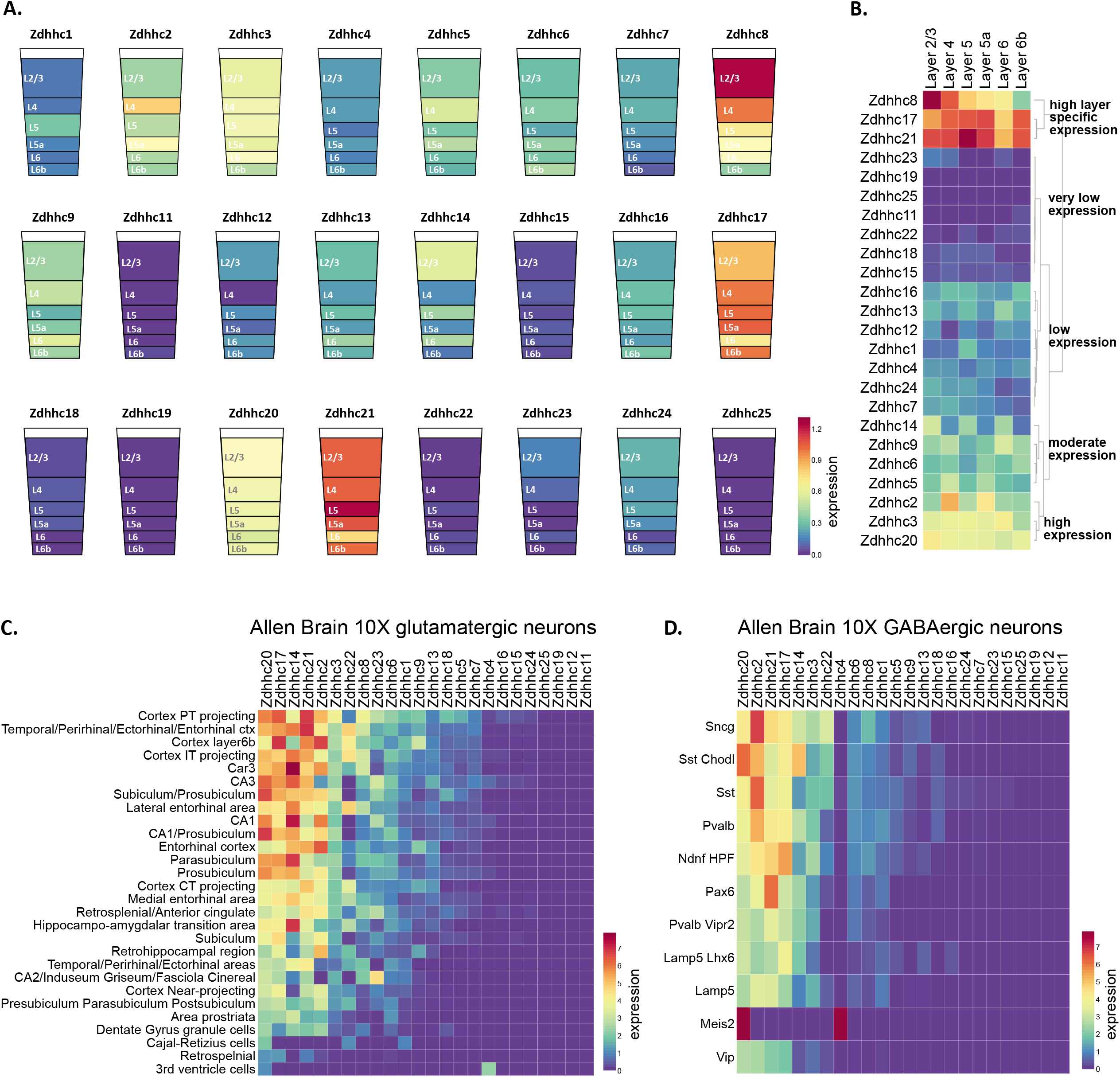
Pyramidal neuron layer specific ZDHHC expression. (A) Heatmap of excitatory neuron ZDHHC expression from somatosensory cortex (original data scRNAseq data from Zeisel et al., 2015) projected onto diagrams of cortical layers. (B) Hierarchical clustering of ZDHHC expression data in A. Heatmap units in (A, B): mean log2(counts per 10,000 + 1) (C) Heatmap of scRNAseq data from Allen Brain 10X genomics (Yao et al., 2021). Data are represented as mean ZDHHC expression per excitatory neuron subtype, with columns and rows sorted by descending mean ZDHHC expression per row/column. (D) As in (C) but for inhibitory neuron subtypes. Heatmap units for (C, D): trimmed mean (25%-75%) Log2(CPM+1)

We examined expression patterns of the ZDHHC enzymes from one of the largest neuronal scRNAseq studies from the isocortex performed by the Allen Brain Institute, which identified 236 glutamatergic and 117 GABAergic distinct neuron metacell types (Yao et al., 2021). We averaged ZDHHC expression data downloaded from BrainPalmSeq for the major metacell clusters from all regions of the isocortex, according to their anatomical location and/or axon projection and plotted ranked heatmaps (Figure 3C,D). We again found that *Zdhhc20* was the ZDHHC transcript with the highest expression, with broad expression across the majority of glutamatergic and GABAergic cell-types. Elevated expression of *Zdhhc14* was found in both glutamatergic and GABAergic neurons, which was moderately expressed in other studies of the brain and cortex discussed previously. Numerous subtypes of projection neurons featured at the top of the ranking including pyramidal tract (PT) projecting neurons found in layer 5 of the cortex, that have extensive dendritic branching and long-range axonal projections to the spinal cord, brainstem and midbrain, as well as the ipsilateral cortex, striatum and thalamus. Other neuron subtypes with high ZDHHC expression included a number of intratelencephalic (IT) projecting neuron classes, including those from Layer 4/5 of the temporal/perirhinal/ectorhinal/entorhinal cortices, Car3 expressing Layer 6 neurons, Layer 6b neurons, and Cortex IT projecting neurons from all cortical layers (Harris and Shepherd, 2015; Yao et al., 2021). GABAergic neurons of the isocortex also showed elevated expression of the common neuronal ZDHHCs including *Zdhhc2*, *Zdhhc3*, *Zdhhc14*, *Zdhhc17*, *Zdhhc20* and *Zdhhc21*. The highest mean ranked expression was observed in the recently categorized Sncg neurons that correspond to *Vip*^+^/*Cck*^+^ multipolar or basket cells (Tasic et al., 2018), and lowest expression was observed in the Vip subclass of interneurons.

Together, our observations reveal that the complex transcriptional diversity of neurons that has recently been revealed by RNA sequencing also includes heterogeneity in the expression of the ZDHHC enzymes that mediate palmitoylation. These expression patterns are likely to influence enzyme-substrate interactions along with the function of *S*-palmitoylation substrates, and thus influence neuronal development, function and synaptic plasticity.

### De-palmitoylating enzyme and ZDHHC accessory protein expression in the nervous system shows regional and cell-type specific patterning

Dynamic turnover of protein *S*-palmitoylation is mediated by the activity of de-palmitoylating enzymes, which determine the half-life of *S*-palmitoylation on a target protein. These include acyl-protein thioesterases 1 and 2 (APT1, APT2; encoded by *Lypla1*, *Lypla2*), palmitoyl-protein thioesterase 1 (PPT1) and the more recently identified α/β hydrolase domain-containing 17 proteins (ABHD17A, ABHD17B and ABHD17C). Compared with the ZDHHC enzymes, relatively less is known about the substrates, sub-cellular localization and brain expression patterns of this family of enzymes. We next explored BrainPalmSeq to determine which cell-types/brain regions show the highest expression of de-palmitoylating enzymes.

We first examined expression heatmaps for the known de-palmitoylating enzymes across the 265 cell-types identified in the ‘MouseBrain’ dataset (Figure 4A) and the cell-type averages for the 565 cell-types from the ‘DropViz’ dataset (Figure 4-figure supplement 1A). *Ppt1* and *Abhd17a* were the enzymes with the broadest expression across all cell-types, with *Ppt1* expression being notably elevated in neurons of the hindbrain and immune cells, and *Abhd17a* being elevated in hindbrain neurons, sensory neurons, oligodendrocytes and epithelial cells. Ranked expression (sorted by descending averages) of de-palmitoylating enzymes in the ‘MouseBrain’ (Figure 4B) and ‘DropViz’ (Figure 4-figure supplement 1B) datasets classified oligodendrocyte lineage cells among the cell-types with the highest expression of de-palmitoylating enzymes overall, primarily due to elevated expression of *Abhd17b*. This mirrors oligodendrocyte enrichment of ZDHHC enzyme expression, again indicating that dynamic regulation of *S*-palmitoylation may be particularly important in this cell type. *Lypla2* expression was greater than *Lypla1* overall in the brain, with *Lypla2* expression being highest in neurons and ependymal cells (Figure 4-figure supplement 1A). *Abhd17c* had the lowest brain expression of all the de-palmitoylating enzymes studied. Correlation analysis between the ZDHHCs and de-palmitoylating enzymes revealed numerous instances of co-expression with almost every ZDHHC (Figure 4D), revealing potential cooperative pairs of palmitoylating and de-palmitoylating enzymes in the nervous system.

**Figure 4.**
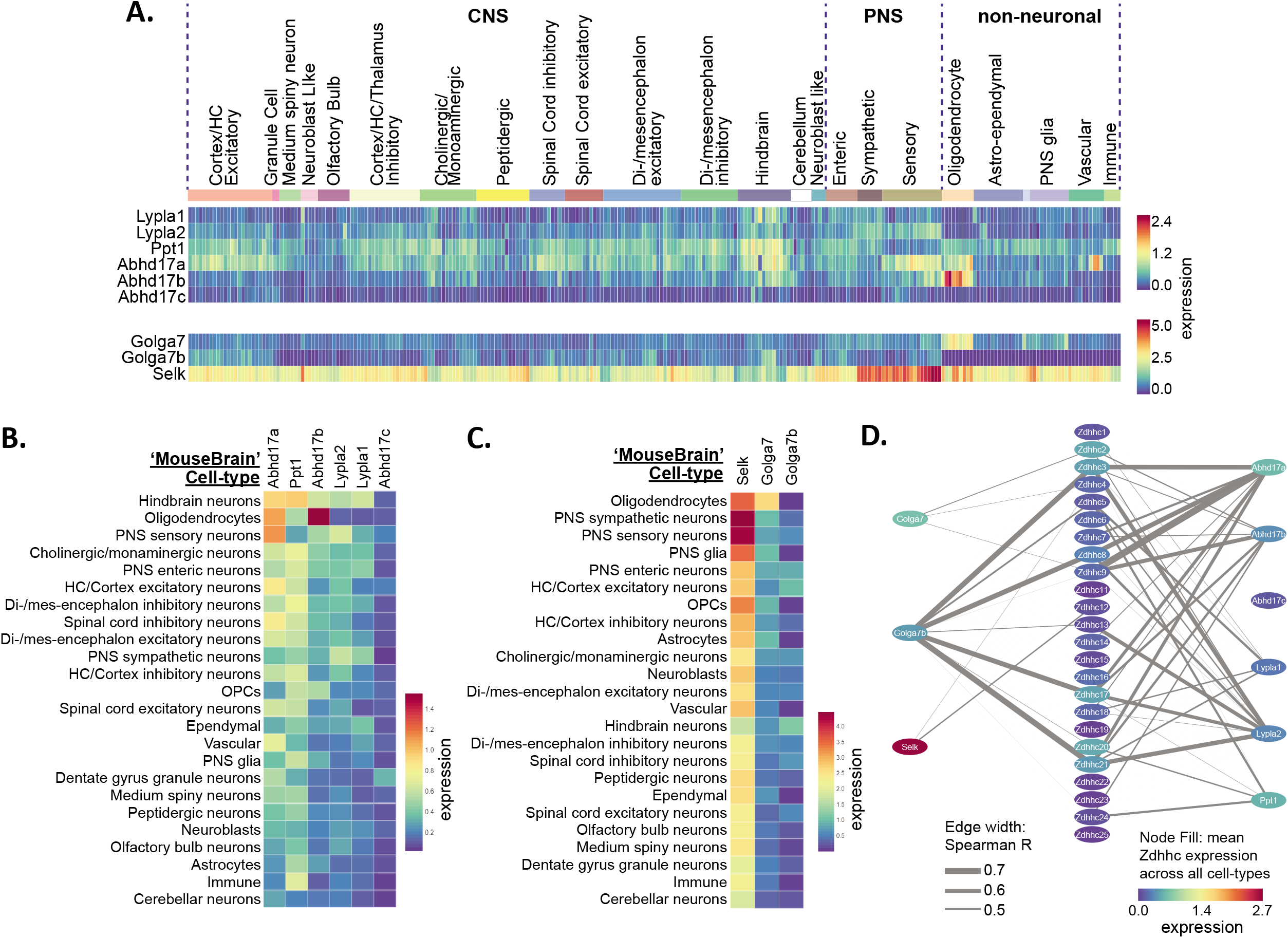
Heterogeneous de-palmitoylating enzyme and ZDHHC accessory protein expression in the mouse nervous system. (A) Heatmap showing expression of de-palmitoylating enzymes (top) and ZDHHC accessory subunits (bottom), extracted from scRNAseq study of mouse CNS and PNS (Zeisel et al., 2018). Each column represents one of the 265 metacells classified in the study. Metacells are organized according to hierarchical clustering designations generated by Zeisel et al. (B) Heatmap showing mean de-palmitoylating enzyme expression per hierarchical cluster, with columns and rows sorted by descending mean ZDHHC expression per row/column. (C) As B but for ZDHHC accessory proteins. (D) Correlation network showing ZDHHC co-expression with de-palmitoylating enyzmes and accessory proteins across all metacells in ‘MouseBrain’ (Spearman R > 0.4). Node color represents mean expression across all metacells. Edge thickness represents strength of correlation.

Although the ZDHHC enzymes are thought to act autonomously, several accessory proteins have been discovered that can regulate ZDHHC stability, localization and catalytic activity (Salaun et al., 2020). These include GOLGA7 (GCP16), which can bind to ZDHHC9 and enhance both protein stability and enzymatic activity by stabilizing the ZDHHC9 auto-palmitoylated intermediate that is formed prior to palmitate transfer from the enzyme to the substrate protein (Mitchell et al., 2014; Swarthout et al., 2005). Both GOLGA7 and related isoform GOLGA7B are also able to interact with ZDHHC5, with the latter influencing ZDHHC5 plasma membrane localization (Woodley and Collins, 2019). Finally, SELK (SELENOK; Selenoprotein K) is an ER localized protein that was found to interact with ZDHHC6, stabilize the auto-palmitoylated intermediate and increase palmitoylation of substrate proteins including the IP_3_ receptor (Fredericks et al., 2017, 2014). We observed widespread expression of *Selk* across all cell-types, with expression being considerably higher than any of the ZDHHCs, de-palmitoylating enzymes or other accessory proteins (Figure 4A, S4A). This is consistent with the known functions of SELK in the ER associated protein degradation pathway and regulation of ER calcium flux (Pitts and Hoffmann, 2018). *Golga7b* expression was widespread across neuronal subtypes but barely detected in glial cells (Figure 4A, S4A). Accordingly, *Golga7b* expression was also strongly correlated with several of the ZDHHCs that were most highly expressed in neurons, including *Zdhhc3*, *Zdhhc8*, *Zdhhc17* and *Zdhhc21* (Figure 4D). In contrast, *Golga7* was enriched in glial cells, particularly in oligodendrocytes, similar to ZDHHC9 for which GOLGA7 is a key accessory protein (Figure 4A, S4A). GOLGA7 and GOLGA7B share 61% amino acid similarity, but their expression was either not correlated or was negatively correlated, indicating that the ZDHHC association of each of these proteins may be regulated in part by differential expression.

### Loss of function mutations in palmitoylating and de-palmitoylating enzymes

Impaired regulation of *S*-palmitoylation has been implicated in numerous neurological disorders, many of which are due to loss of function (LOF) mutations in the genes encoding palmitoylating and de-palmitoylating enzymes (Cho and Park, 2016; Matt et al., 2019). We next sought to determine if the regional and cell type expression data available in BrainPalmSeq could reveal insights into the pathogenesis of disorders caused by LOF mutations in palmitoylating and de-palmitoylating enzymes. As many of these diseases have a neurodevelopmental origin, we examined whole brain datasets curated in BrainPalmSeq from the neonatal (Rosenberg et al., 2018), adolescent (Zeisel et al., 2018) and adult (Sjöstedt et al., 2020) mouse brain.

A single nucleotide polymorphism (SNP) in the *ZDHHC8* gene has been implicated in increased susceptibility to schizophrenia (Chen et al., 2004; Mukai et al., 2004), while hemizygous microdeletion in the chromosomal locus 22q11, which encodes a number of genes including *ZDHHC8*, is one of the highest known genetic risk factors to developing schizophrenia (Figure 5A; Karayiorgou et al., 2010). To assess the developmental expression of *Zdhhc8*, we averaged expression within broadly defined cell-type clusters that could be applied to both the Rosenberg and Zeisel scRNAseq datasets (Figure 5A, B; Supplementary File 3). *Zdhhc8* expression was highest in neurons of the cortex and hippocampus, followed by neurons of the mid-and hindbrain at both developmental ages. To explore regional expression in the adult mouse brain, we projected BrainPalmSeq generated heatmaps expression data from the ‘Protein Atlas’ mouse whole brain dataset (bulk RNAseq from major brain regions; Figure 5-figure supplement 1) onto anatomical maps of the mouse brain, again revealing highest expression of *Zdhhc8* in the cortex, followed by the hippocampus and basal ganglia (Figure 5C; Sjöstedt et al., 2020). *Zdhhc8* expression was particularly enriched in Layer 2/3 of the neonatal (not shown) and adult mouse (Figure S3B) cortex, which is the cortical layer with the most pronounced morphological deficits in patients with Schizophrenia (Glantz and Lewis, 2000; Kolluri et al., 2005; Wagstyl et al., 2016). Together, we found *Zdhhc8* expression patterns in the mouse brain that are established early in postnatal development and maintained into adulthood, that also overlay with many brain regions and cell types that are known to be severely affected in patients with schizophrenia. These observations support a model in which LOF *ZDHHC8* mutations may elicit many of the symptoms of schizophrenia by disrupting *S*-palmitoylation and normal neuronal development in these brain regions.

**Figure 5.**
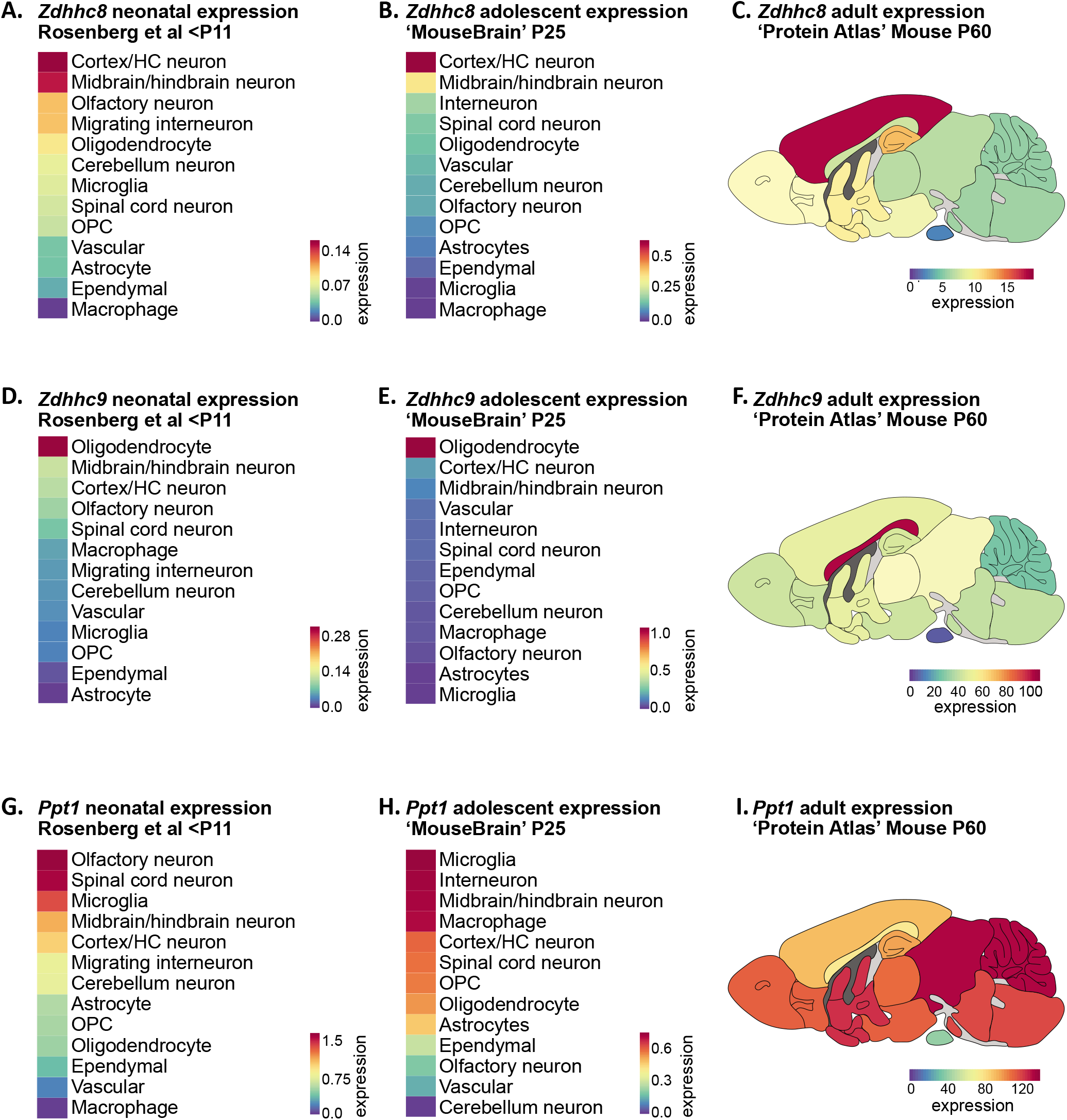
Disease associated palmitoylating enzyme regional and cell-type expression overlays with brain pathology in associated LOF disorders. (A) Heatmap showing ranked *Zdhhc8* expressing neuronal and glial cell types in descending order. Original data from scRNAseq neonatal mouse brain study; Rosenberg et al., 2018. Cell types were averaged as described in Supplementary File 3. Heatmap units: mean log2(counts per 10,000 + 1). (B) As in A, but original data from scRNAseq adolescent mouse brain study Zeisel et al (2018). Heatmap units: mean log2(counts per 10,000 + 1). (C) Heatmap of *Zdhhc8* expression from whole brain regional bulk RNAseq data (original data from ‘Protein Atlas’; Sjöstedt et al., 2020) projected onto anatomical map of mouse brain. Heatmap units: pTPM. (D-F) As in (A)-(C) but for *Zdhhc9*. (G-I) As in (A)-(C) but for *Ppt1*.

Mutations in the *ZDHHC9* gene, which is located on the X chromosome, have been identified in ∼2% of individuals with X-linked intellectual disability (ID) (Raymond et al., 2007; Tzschach et al., 2015). Neuroanatomical abnormalities reported in patients with *ZDHHC9* mutations include decreased cortical, thalamic and striatal volume, as well as widespread white matter abnormalities with prominent hypoplasia (under-development) of the corpus callosum (Baker et al., 2015; Bathelt et al., 2016). Disrupted white matter integrity is thought to underlie deficits in global and local brain connectivity in patients with *ZDHHC9* mutations (Bathelt et al., 2017). *Zdhhc9* knock-out mice also develop similar pathological changes, including decreased volume of the corpus callosum (Kouskou et al., 2018). We observed considerable cell-type enrichment of *Zdhhc9* in oligodendrocytes across studies and developmental ages (Figure 5D, E), accompanied by moderate neuronal expression of *Zdhhc9* relative to other ZDHHCs across several brain regions including the hippocampus and cortex (Figure 2A, 3A, 5D, E), consistent with the known function of ZDHHC9 in regulating neuronal development (Shimell et al., 2019). Regionally, we found *Zdhhc9* expression in adult mice to be highly enriched in the corpus callosum, the largest white matter tract in the brain (Figure 5F). As myelin production by oligodendrocytes is critical for maintaining white matter integrity, these observations indicate that disrupting *S*-palmitoylation in oligodendrocytes may underlie the white matter pathology and decreased connectivity observed in patients with X-linked ID and *ZDHHC9* mutations.

Infantile neuronal ceroid lipofuscinosis (INCL or CLN1 disease) is a severe neurological disorder caused by LOF mutations in the *PPT1* gene that presents in the first 6-12 months of life and is characterized by rapid developmental regression, blindness and seizures, with continual deterioration until death in early childhood (Nita et al., 2016). While PPT1 is thought to primarily localize to lysosomes with an essential role in lysosomal degradation of *S*-palmitoylated proteins (Lu et al., 1996), this protein also has a synapse-specific function in regulating synaptic vesicle cycling and synaptic transmission (Koster and Yoshii, 2019). We found that neuronal *Ppt1* expression was high in postnatal neurons of the spinal cord, olfactory bulb and mid/hindbrain, while microglia were the highest expressing non-neuronal cell type at both postnatal ages (Figure 5G, H). Neurodegeneration has been detected in the spinal cord prior to onset within the brain in *Ppt1* knock-out mice, accompanied by extensive glial cell activation including microgliosis, which is a pathological hallmark of CLN1 disease (Shyng et al., 2017). Mid-/hindbrain neurons also had high expression of *Ppt1*, consistent with reports that *Ppt1* knock-out mice show early signs of brain pathology in the thalamus (Kielar et al., 2007). Overall, we observed widespread *Ppt1* expression in almost every brain region in adult mice, consistent with the sweeping neurological deficits associated with CLN1 disease (Figure 5I). Together, these observations reveal how the loss of *Ppt1* in cell types with high *Ppt1* expression may lead to cell death/dysfuntion in the early stages of CLN1 disease.

### ZDHHC cell type enrichments can be used to predict and validate ZDHHC substrates

We next tested if ZDHHC expression patterns identified from BrainPalmSeq could be used to predict and validate *S*-palmitoylation substrates for regionally enriched ZDHHCs. We focused on *Zdhhc9*, which showed a consistent cell-type enrichment in oligodendrocytes across multiple studies in BrainPalmSeq, while LOF mutations in *ZDHHC9* are known to result in reduced white matter integrity in the brain (Raymond et al., 2007). Examination of the Marques et al oligodendrocyte-specific scRNAseq dataset curated in BrainPalmSeq revealed that oligodendrocyte *Zdhhc9* expression increased throughout maturation, with highest expression in the myelin forming (MFOL) intermediate-maturity subtype of oligodendrocytes, while slightly lower expression is maintained in mature oligodendrocytes (MOL; Figure 6A; Marques et al., 2016).

**Figure 6.**
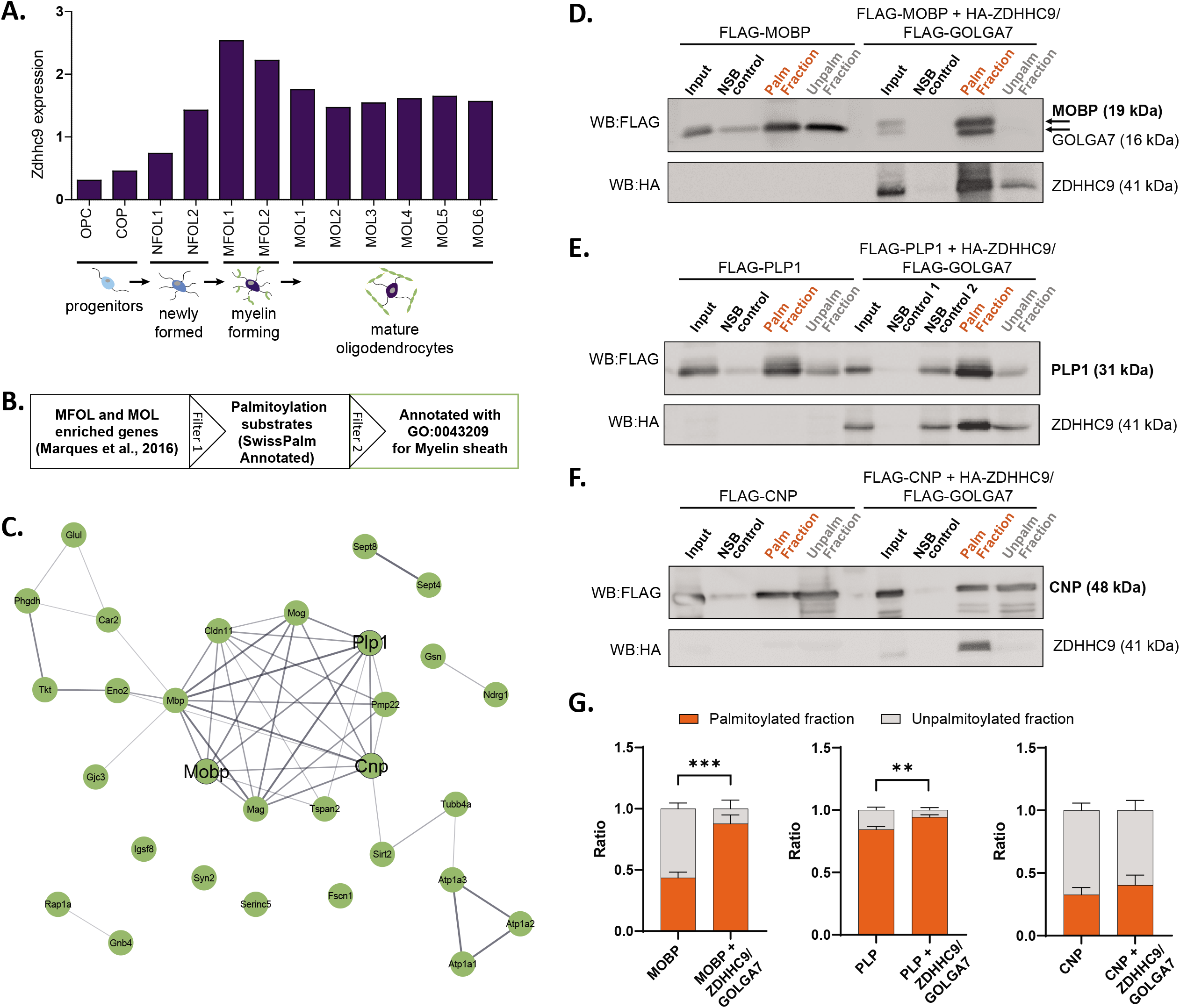
Validation of predicted *S*-palmitoylation substrates of *Zdhhc9* derived from cell-type enriched expression. (A) Graph of expression data for *Zdhhc9* extracted from BrainPalmSeq. Original data from oligodendrocyte scRNAseq study by Marques et al. Expression units: mean log2(counts per 10,000 + 1). (B) Diagram illustrating workflow to generate a list of oligodendrocyte enriched palmitoylation substrates, GO annotated for myelin sheath for experimental validation. (C) STRING diagram of myelin sheath annotated palmitoylation substrates. (D) Western blot following Acyl-RAC palmitoylation assay in HEK293 cells to identify palmitoylated and unpalmitoylated fractions of FLAG-MOBP either without or with co-transfection of FLAG-GOLGA7 and HA-ZDHHC9. Input = unprocessed protein lysate. NSB control = non-specific binding of unpalmitoylated protein to sepharose resin in control pipeline. Palm fraction = palmitoylated protein (bound to sepharose resin). Unpalm fraction = unpalmitoylated protein (did not bind to sepharose resin). (E-F) As in (D) but for FLAG-PLP1 (E) or FLAG-CNP (F). (G) Graphs quantifying the ratio of palmitoylated to unpalmitoylated protein either with or without co-transfections with FLAG-GOLGA7 and HA-ZDHHC9. *n* = 4-6 HEK cell cultures per condition. Statistics shown for palmitoylated fraction. Two-way ANOVA; Šídák’s *post hoc*; mean ± SEM. MOBP: p = 0.0004, 95% CI −0.6594 to −0.2266; PLP1: p = 0.0046, 95% CI −0.1660 to −0.03011; CNP: p = 0.6981, 95% CI - 0.3274 to 0.1742.

To identify potential substrates for ZDHHC9, we cross-referenced a list of MFOL/MOL enriched genes identified in the study by Marques et al (Marques et al., 2016) with the SwissPalm database to identify known palmitoylation substrates in these cell types (Swiss Palm Annotated; Figure 6B; Blanc et al., 2019, 2015). PANTHER analysis of cellular component enrichments for these substrates revealed the most significant enrichment was for the term ‘myelin sheath’ (30 proteins; Figure 6C, Figure 6-figure supplement 1). To determine if any of the myelin sheath associated proteins could be palmitoylated by ZDHHC9, we selected three proteins (MOBP, PLP1 and CNP) for experimental validation (Figure 6C). We separately expressed each of these candidate substrates together with ZDHHC9 and its accessory protein GOLGA7 in HEK293T cells, and determined the proportion of palmitoylated substrate using an acyl-resin assisted capture (acyl-RAC) palmitoylation assay (Forrester et al., 2011). In this assay, free cysteine residues of cell lysates were first blocked, followed by cleavage of the palmitoyl-thioester bond with hydroxylamine, resulting in the exposure of a free sulfhydryl group. Cleaved lysates were then applied to a Sepharose resin to capture palmitoylated proteins containing a free sulfhydryl group. Un-bound proteins (Unpalm Fraction) were first extracted from the resin mixture, followed by elution of bound palmitoylated proteins from the resin (Palm Fraction). To assess non-specific binding of unpalmitoylated protein to the resin, half of the cell lysate was processed without the cleavage step (NSB control). Co-expression of HA-ZDHHC9 and FLAG-GOLGA7 increased the palmitoylated fraction of MOBP and PLP1, indicating that these proteins are substrates for ZDHHC9 (Figure 6D, E, G). Conversely, CNP was not identified as a ZDHHC9 substrate in our assay (Figure 6F, G). These results demonstrate how the cell-type enrichments of ZDHHC enzymes identified in this study can be used, along with the lists of similarly enriched palmitoylation substrates, to guide the identification of enzyme-substrate interactions that can be further investigated *in vivo*.

## Discussion

### BrainPalmSeq as a tool to generate hypotheses about proteins that control S-palmitoylation in the brain

We have demonstrated the utility of BrainPalmSeq by providing examples of how this database can be used to explore detailed region and cell type-specific expression patterns of the known palmitoylating and de-palmitoylating enzymes, and their accessory proteins. We reveal how these expression patterns can be used to predict/validate *S*-palmitoylation substrates and better understand diseases associated with loss of function mutations in the enzymes that mediate *S*-palmitoylation. Given the number of brain regions and cell types incorporated into BrainPalmSeq that were not discussed in the present study, including the thalamus, hypothalamus, amygdala, striatum and cerebellum, there is rich potential for users to explore the data and generate hypotheses about the role of these enzymes in the brain.

### Insights into the role of *S*-palmitoylation associated enzymes in brain physiology and pathology

While we found that many of the proteins we studied showed correlated expression across the entire mouse nervous system, particularly those enriched in neurons including *Zdhhc3*, *Zdhhc8*, *Zdhhc17* and *Zdhhc21*, expression of these genes was segregated within more narrowly defined neuronal populations such as the excitatory pyramidal neurons within the hippocampal tri-synaptic loop or layers of the somatosensory cortex. This is in line with the extensive neuronal transcriptional heterogeneity identified recently by a number of scRNAseq studies (Saunders et al., 2018; Yao et al., 2021; Zeisel et al., 2015, 2018). The genes that determine neuronal identity fall under four broad functional categories: those that control transcriptional programs, membrane conductance, neurotransmission, or synaptic connectivity (Zeisel et al., 2018). We also report heterogeneity in the neuronal fingerprint of palmitoylating and de-palmitoylating enzyme expression, which will in turn give rise to differential *S*-palmitoylation of neuronal proteins. Future work is needed to determine how these specific ZDHHC expression patterns are related to dynamic *S*-palmitoylation in these neuronal sub-types, and how the elevated expression of certain ZDHHCs can alter neuronal function. Given that *S*-palmitoylation is a key regulator of neuronal development, and that nearly half of all known synaptic proteins are substrates for palmitoylation (Sanders et al., 2015), this heterogeneity is likely to be a key mechanism in the fine tuning of neuronal function and synaptic transmission.

Many of the ZDHHCs that we observed with consistently elevated expression across multiple studies in BrainPalmSeq have already been studied in the context of neuronal signaling, including ZDHHC2, ZDHHC3, ZDHHC8 and ZDHHC17 (Ji and Skup, 2021; Matt et al., 2019). In contrast, ZDHHC20 and ZDHHC21 are relatively understudied in the nervous system, despite our observation that these are two of the most abundantly expressed ZDHHCs across neuronal cell types, with broad expression of *Zdhhc20* also in glial cells. A recent study defined a role for ZDHHC21 in the palmitoylation of serotonergic receptor 5-HT1A and implicated downregulation of ZDHHC21 in the development of major depressive disorder (Gorinski et al., 2019). Interestingly, both ZDHHC20 and ZDHHC21 have a potential role in the pathogenesis of Alzheimer’s disease, as they can palmitoylate BACE1, Tau and amyloid precursor protein (Cho and Park, 2016). Further work is required to understand the likely important role of these enzymes in the brain.

We made several other interesting observations during our examination of BrainPalmSeq that were not discussed in detail in the present study but we believe warrant further investigation. For example, the particularly elevated expression of *Zdhhc2* in peripheral sensory neurons may indicate an important role for palmitoylation in this cell type. Across multiple studies we observed striking enrichment of *Zdhhc14* in cerebellar Purkinje neurons, a cell type in which *S*-palmitoylation is known to be important for long-term depression, although the role of ZDHHC14 in this process has not yet been investigated (Thomas et al., 2013). *Zdhhc23* was similarly enriched in the CA2 region of the hippocampus, with comparatively low expression across other cell types. More broadly, the elevated expression of a variety of palmitoylating enzymes in neurons that utilize acetylcholine or monoamines as neurotransmitters would suggest an important role for *S*-palmitoylation in these neurons that has yet to be explored. Accordingly, many of the key proteins involved in cholinergic synaptic transmission are *S*-palmitoylation substrates including muscarinic acetylcholine receptor M2 (CHRM2), acetylcholinesterase (ACHE) and ATP-citrate synthase (ACLY; Blanc et al., 2015, 2019). Our observations of co-enrichment of certain palmitoylating and de-palmitoylating enzymes are also of interest, such as *Abhd17b* and *Zdhhc9* in oligodendrocytes. It is possible that these enzymes share substrates to mediate dynamic palmitoylation/de-palmitoylation, or conversely, have separate substrates in order to maintain stable *S*-palmitoylation states of certain oligodendrocyte expressed proteins. Importantly, the data accessibility in BrainPalmSeq will enable researchers to develop hypotheses regarding their cell type, brain region or protein of interest.

The palmitoylome of each cell type in the nervous system is likely to be highly heterogeneous and will be determined by the expression of both the *S*-palmitoylation substrates and the palmitoylating and de-palmitoylating enzymes in a given cell type. Furthermore, accumulating evidence has revealed that this palmitoylome can be altered by extrinsic factors such as chronic stress and neuronal activity (Kang et al., 2008; Zareba-Koziol et al., 2019). While we have provided predicted palmitoylomes composed of several highly expressed or enriched *S*-palmitoylation substrates in select brain regions and cell types, experimental validation to reveal the relative palmitoylation of substrates under various conditions is needed to fully understand these cellular differences. Nevertheless, we were able utilize our predicted palmitoylomes to validate substrates for ZDHHC9, providing insight into the potential role of this enzyme in myelin regulation in the brain.

Neurological disorders that arise from LOF gene mutations may be predicted to lead to pathological changes that are more severe in the brain regions in which these genes are most highly expressed. We observed this type of regional overlay for the expression patterns of *Zdhhc8*, *Zdhhc9* and *Ppt1*. Numerous other brain disorders are thought to be exacerbated by an imbalance in *S*-palmitoylation, such as decreased *S*-palmitoylation of HTT in Huntington’s disease (Virlogeux et al., 2021; Yanai et al., 2006), increased *S*-palmitoylation of APP and TAU in Alzheimer’s disease (Cho and Park, 2016), and reduced *S*-palmitoylation of 5-HTA receptor in major depressive disorder (Gorinski et al., 2019). Efforts are already underway to normalize aberrant *S*-palmitoylation in neurological diseases in order to improve clinical outcomes (Roberts et al., 2012; Virlogeux et al., 2021). Understanding the brain expression patterns of the enzymes that mediate palmitoylation in these diseases will be paramount to developing and targeting such therapeutics.

### Differential gene expression as a means to control *S*-palmitoylation in the brain

The mechanisms that govern ZDHHC enzyme-substrate interactions are complex and still not fully understood. While the majority of post-translational modifications including phosphorylation and N-glycosylation are highly sequence specific (Schwarz and Aebi, 2011; Ubersax and Ferrell, 2007), several studies have revealed that *S*-palmitoylation by ZDHHCs can be stochastic, proximity based and lacking in stereo-selectivity (Rocks et al., 2010; Rodenburg et al., 2017). Contrasting studies have shown that numerous ZDHHCs have specific protein interacting domains including ankyrin repeat (AR), PDZ and SH3 domains that facilitate substrate interactions, providing support for a model in which more specific enzyme-substrate interactions can govern *S*-palmitoylation (Abrami et al., 2017; Lemonidis et al., 2015; Plain et al., 2020; Rana et al., 2018; Thomas et al., 2012; Verardi et al., 2017). Furthermore, a recent study found striking substrate specificity for several ZDHHCs with the G-protein subunit Gαo, and revealed intriguing observations that the subcellular localization of a number of *S*-palmitoylation substrates could be controlled by changing the localization, and importantly, the expression of certain ZDHHC enzymes. In this study, *S*-palmitoylated substrates were found to accumulate in the subcellular compartment in which their partner ZDHHCs were targeted (Solis et al., 2020). This is particularly relevant as the ZDHHCs are known to have diverse subcellular localizations including the Golgi, ER, endosomes and plasma membrane (Globa and Bamji, 2017). Transcriptional control of differentially compartmentalized palmitoylating and de-palmitoylating enzymes could therefore be an essential mechanism for regulating the subcellular localization, and function, of *S*-palmitoylated protein substrates. Accordingly, LOF mutations in certain ZDHHC enzymes leads to cell type-specific disruption in *S*-palmitoylation that is not compensated by other members of the large ZDHHC family. We provide a means to investigate the expression of the proteins that mediate *S*-palmitoylation, making BrainPalmSeq an invaluable resource to both researchers and clinicians that are working to better understand the role of *S*-palmitoylation in the brain.

## Source Data Legends

**Figure 1 – Source data 1.** ZDHHC expression in the mouse nervous system

**Figure 2 – Source data 1.** ZDHHC expression and S-palmitoylation substrate expression in the hippocampus

**Figure 3 – Source data 1.** ZDHHC expression cortical neurons

**Figure 4 – Source data 1.** De-palmitoylating enzyme and ZDHHC accessory protein expression in the mouse nervous system

**Figure 5 – Source data 1.** Disease associated palmitoylating enzyme regional and cell-type expression patterns

**Figure 6 – Source data 1.** Validation of predicted *S*-palmitoylation substrates of *Zdhhc9*

## Materials and Methods

### Data processing for BrainPalmSeq

For Zeisel et al., 2018 (‘MouseBrain’), single-cell counts (UMI from 3’ end counting) were downloaded from MouseBrain.org (loom file named l5_all.loom), and log normalized by first scaling the expression values provided to a sum of 10,000 per cell before calculating log2(scaled_counts+1). Averages were then performed by brain region, neurotransmitter and taxonomy for each gene.

For DropVIz Metacell counts were downloaded from DropViz.org (count file metacells.BrainCellAtlas_Saunders_version_2018.04.01.RDS and annotation file annotation.BrainCellAtlas_Saunders_version_2018.04.01.RDS) and log normalized by first scaling the expression values provided to a sum of 10,000 per metacell before calculating log2(scaled_counts+1). Averages were then performed by cell type, tissue and class for each gene. Genes associated with palmitoylation were selected in order to create the heatmaps.

For Zeisel, single-cell counts (UMI from 3’ end counting) were downloaded from https://storage.googleapis.com/linnarsson-lab-www-blobs/blobs/cortex/expression_mRNA_17-Aug-2014.txt, and log normalized by first scaling the expression values provided to a sum of 10,000 per cell before calculating log2(scaled_counts+1). Averages were then performed by cluster, tissue and class for each gene. Genes associated with palmitoylation were selected in order to create the heatmaps, categories comprising fewer than 5 single cells are not displayed.

For Marques, single-cell counts (UMI from 3’ end counting) were downloaded from GEO with accession ID GSE75330 (file GSE75330_Marques_et_al_mol_counts2.tab) and log normalized by first scaling the expression values provided to a sum of 10,000 per cell before calculating log2(scaled_counts+1). Averages were then performed by cluster and region for each gene. Genes associated with palmitoylation were selected in order to create the heatmaps, categories comprising fewer than 5 single cells are not displayed.

For Rosenberg, single-cell counts (UMI from 3’ end counting) were downloaded from GSE110823, and log normalized by first scaling the expression values provided to a sum of 10,000 per cell before calculating log2(scaled_counts+1). Averages were then performed by brain region, neurotransmitter and taxonomy for each gene. Genes associated with palmitoylation were selected in order to create the heatmaps.

Data from Sjöstedt el al. were downloaded as Protein-coding transcripts per million (pTPM) from proteinatlas.org (“RNA mouse brain region gene data”) and not further processed.

For Hipposeq, expression data were downloaded as FKPM directly from hipposeq.janelia.org and were not further processed.

For Allen Brain 10X data, expression data were downloaded as trimmed means (25%-75%) Log2(CPM+1) from portal.brain-map.org/ and were not further processed.

### Correlation analysis

Spearman correlation values between genes and their significances were calculated in R using the expression results obtained for each cell type as described above.

### Identification of *S*-palmitoylation substrates with SwissPalm

Gene lists were inputted into SwissPalm (https://swisspalm.org/proteins) input file function and cross-referenced with ‘Dataset 3: Palmitoylation validated or found in at least one palmitoyl-proteome (SwissPalm annotated)’ for *Mus musculus*, with an additional filter for UniProt ‘Reviewed’ proteins.

### Generating a predicted palmitoylome for dorsal hippocampus

To curate the regionally enriched predicted-palmitoylome, the enrichment analysis tools in hipposeq (https://hipposeq.janelia.org/) were used to compare each of the selected Cell Lines vs the other Cell Lines in the analysis (Selected Cell Lines = dorsal DG, CA3, CA2 and CA1), with the following parameters: ‘Fold threshold’ = 1.5; ‘FKPMmin threshold’ = 5, ‘FDR’ = 0.05. The resulting lists of regionally enriched transcripts were cross referenced with SwissPalm as described above to identify regionally enriched *S*-palmitoylation substrates.

### Bioinformatic analysis

Gene Ontology (GO) analysis was performed using statistical overrepresentation tests in PANTHER16.0 (Mi et al., 2009) with default settings and *Mus musculus* as the reference species. Biological process GO terms were extracted and ranked according to false discovery rate (FDR). Kyoto Encyclopedia of Genes and Genomes (KEGG) analysis was performed using the web-based program Enrichr (Chen et al., 2013; Kuleshov et al., 2016) and ranked according to −log Adjusted P-value. Synaptic Gene Ontologies (SynGO; version 1.1) analysis was performed using default settings with brain expressed genes as a background and terms for ‘biological process’ were ranked according to −log Adjusted P-value. Functional protein interaction networks were identified using the Search Tool for the Retrieval of Interacting Genes (STRING) 11.0 (Szklarczyk et al., 2019) with *Mus musculus* as the reference species. Seven types of protein interactions were used for network generation, including text mining, neighborhood, co-occurrence, co-expression, gene fusion, experiments and databases.

### Data presentation

Heatmaps were plotted and hierarchical clustering performed in Displayr (https://www.displayr.com) using the ‘Dendrogram’ function. Cytoscape (Version 3.8.0) was used to draw correlation networks.

### Heatmap creation for BrainPalmSeq

All plots for the BrainPalmSeq database were generated using curated RNA sequencing datasets. Python 3 and Javascript scripts were used with the plotting library Bokeh to generate the interactive heatmaps to display and compare these datasets on the BrainPalmSeq website (Bokeh Development Team, 2018).

### Cell culture

HEK293T cells were thawed and aliquoted into a 10cm dish with 10mL prewarmed (37°C) DMEM (GIBCO, Thermo Fisher Scientific, Waltham, MA) supplemented with 10% fetal bovine serum (FBS) (GIBCO, Thermo Fisher Scientific, Waltham, MA) and 1% Pen/Strep(P/S) (GIBCO, Thermo Fisher Scientific, Waltham, MA). HEK293T cells were then placed in a 37°C incubator with 5% CO2 and passaged approximately every 5 days, or once confluency was achieved.

### Transfection

70%-80% confluent HEK293T cells were transfected using Lipofectamine 2000 (Invitrogen/Life Technologies, Carlsbad, CA) according to the manufacturer’s recommendations. Each well of a 6-well plate was transfected with a total of 3ug DNA, 150uL of Opti-Mem (GIBCO, Thermo Fisher Scientific, Waltham, MA) was used with 6uL of Lipofectamine 2000 (Invitrogen/Life Technologies, Carlsbad, CA). Experimental condition wells were transfected with 1ug of the indicated construct of interest, 1ug of HA-DHHC9 (mouse; Shimell et al., 2019), and 1ug of FLAG-GOLGA7 (Maurine Linder, Washington University School of Medicine). Human FLAG-MOBP (CAT#: RC223946), FLAG-PLP1 (CAT#: RC218616) and FLAG-CNP (CAT#: RC207038) were acquired from Origene, Maryland, USA. Control condition wells were transfected with 1ug of the indicated construct of interest, and 2ug of a scrambled control plasmid. Cells were lysed using the acyl-RAC assay lysis buffer 48hours after transfection.

### Palmitoylation Assay (acyl-RAC)

The commercially available CAPTUREome S-palmitoylated protein kit (Badrilla, Leeds, UK) was used in accordance with the manufacturer’s guidelines with three optimizations: (1) prior to the cell lysis step, wells were washed with 1mL of 1X PBS to eliminate any dead cells or residual media; (2) during the cell lysis step, DNase (Sigma-Aldrich, St. Louis, MO), was added to the solution (5uL per 500uL of lysis buffer); and (2) protein concentration was measured prior to the separation of experimental sample and negative control sample using the BCA Assay (Pierce, Thermo Fisher Scientific, Waltham, MA).

### Western Blot Analysis

Western blotting was performed using 4% stacking and 12% resolving SDS-PAGE gels. PVDF membranes were then blocked for 1hour at room temperature with 5% BSA in 0.05% TBS-T. PVDF membranes were incubated with the indicated primary antibodies (anti-HA: Cell Signaling Technology, C29F4, Rabbit mAb CAT#: 3724, 1:1000; anti-FLAG: Origene, mouse monoclonal antibody, CAT#: TA50011-100, 1:1000) overnight at 4°C. Proteins were then visualized using enhanced chemiluminescence (Immubilon Western Chemiluminescent HRP Substrate) on a BioRad ChemiDoc XRS+ scanner. Blots were then quantified using Fiji1 software. The palmitoylated and unpalmitoylated fractions were calculated using the following equations respectively: (Palm Fraction / (Palm Fraction + Unpalm Fraction)) and (Unpalm Fraction / (Palm Fraction + Unpalm Fraction)). Criteria for data inclusion were sufficient transfection/antibody signal and minimal protein in the non-specific binding control.

### Statistical Analysis

Spearman correlations were performed with the function cor in R, and their significances were obtained using the function cor.test followed by a correction for multiple testing using p.adjust with the fdr method (Benjamini & Hochberg). For assessing enrichment of *S*-palmitoylation substrates within Zdhhc co-expressed genes, Fisher’s exact test was performed using the function fisher.test in R.

For validation of substrates using Acyl-Rac palmitoylation assay, two-way ANOVA with Šídák’s *post hoc* was used to assess significance of both palmitoylated and unpalmitoylated fractions. Statistics are shown for palmitoylated fraction. No outliers were excluded. Statistical analyses were performed in GraphPad Prism 9.2.0 (San Diego, CA, USA). *n* represents the number of individual HEK cell culture dishes. Each culture dish is defined as a biological replicate. No technical replicates were performed.

## Acknowledgements

The authors would like to thank Drs. A. Ciernia, M. Cembrowski, T. Murphy and J. LeDue for helpful discussion. This work was supported by CIHR Foundation grant (F18-00650) to SXB and by computational resources made available through the NeuroImaging and NeuroComputation Centre at the Djavad Mowafaghian Centre for Brain Health (RRID SCR_019086) and the Dynamic Brain Circuits in Health and Disease Research Excellence Cluster DataBinge Forum.

