## Supplemental Figs for "BrainPalmSeq: A curated RNA-seq database of palmitoylating and de-palmitoylating enzyme expression in the mouse brain"

Figure 1-figure supplement 1

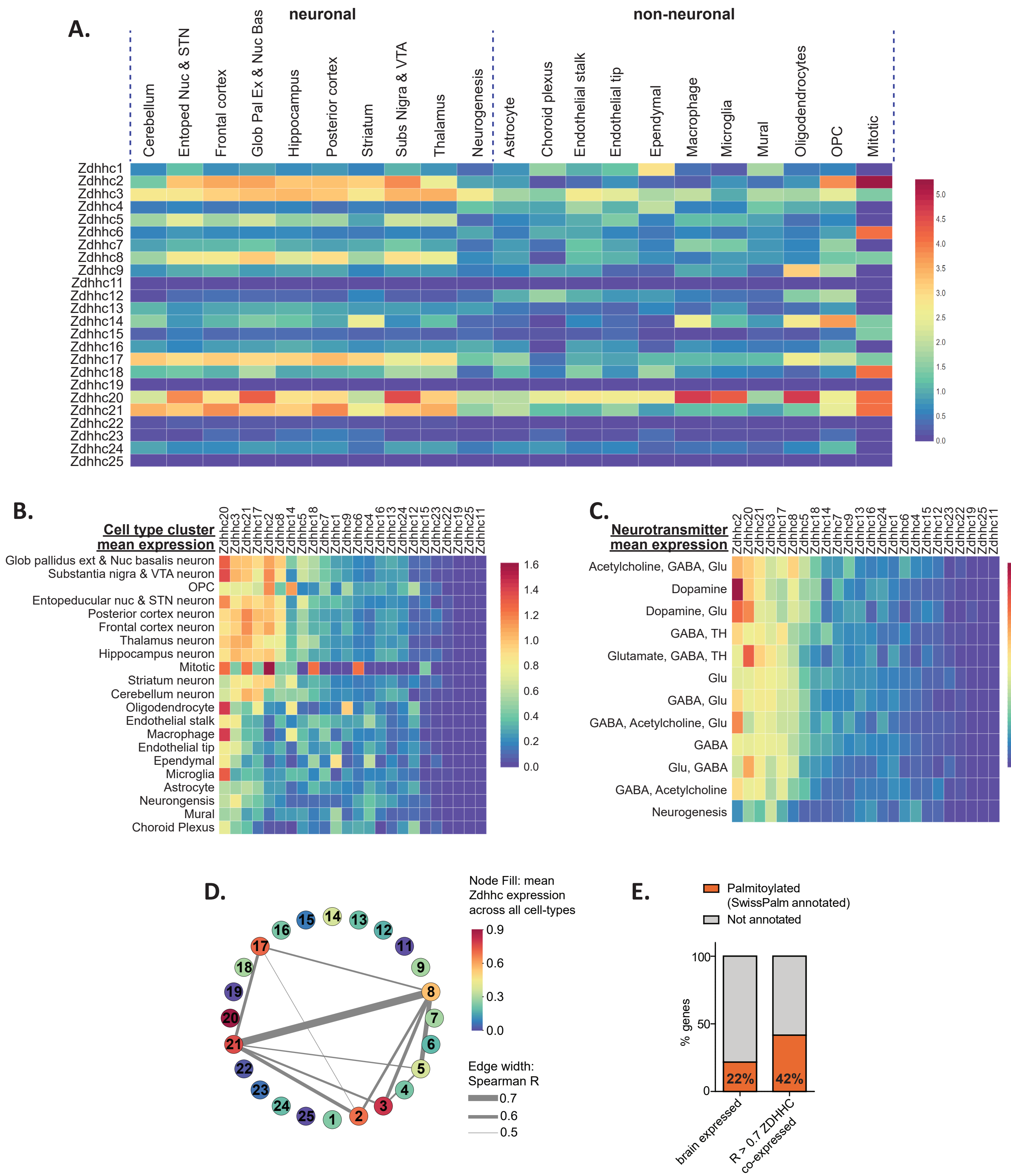

**Figure 1-figure supplement 1. Heterogeneous ZDHHC expression in the mouse brain**

(A) Heatmap showing expression for the 24 ZDHHC genes, extracted from 'DropViz' scRNAseq study of mouse brain (Saunders et al., 2018). Each column represents metacell averages. Metacells are organized along x-axis according to brain region (neuronal) or class (non-neuronal). Full metadata for this study available on BrainPalmSeq.

(B) Heatmap showing mean Zdhhc expression per brain region/class, with columns and rows sorted by descending mean Zdhhc expression per row/column.

(C) Heatmap showing mean Zdhhc expression per neurotransmitter cluster for neurons from all brain regions. Columns and rows are sorted as in B.

(D) Correlation network showing ZDHHC co-expression across all metacells in 'DropViz' (Spearman R > 0.5). Numbers in nodes correspond to ZDHHC number. Node color represents mean expression across all metacells. Edge thickness represents strength of correlation.

(E) Graph showing proportion of genes from 'DropViz' dataset that are co-expressed with one or more ZDHHC and also substrates for S-palmitoylation (SwissPalm annotated). 'Brain expressed' = 15,389 protein coding genes expressed in the postnatal mouse brain, curated from the MGI RNAseq studies database. 'R > 0.7 ZDHHC co-expressed' = 676 genes co-expressed with one or more ZDHHC (Spearman R > 0.7).

Units for all heatmaps in figure: mean log2(counts per 10,000 + 1).

Figure 2-figure supplement 1

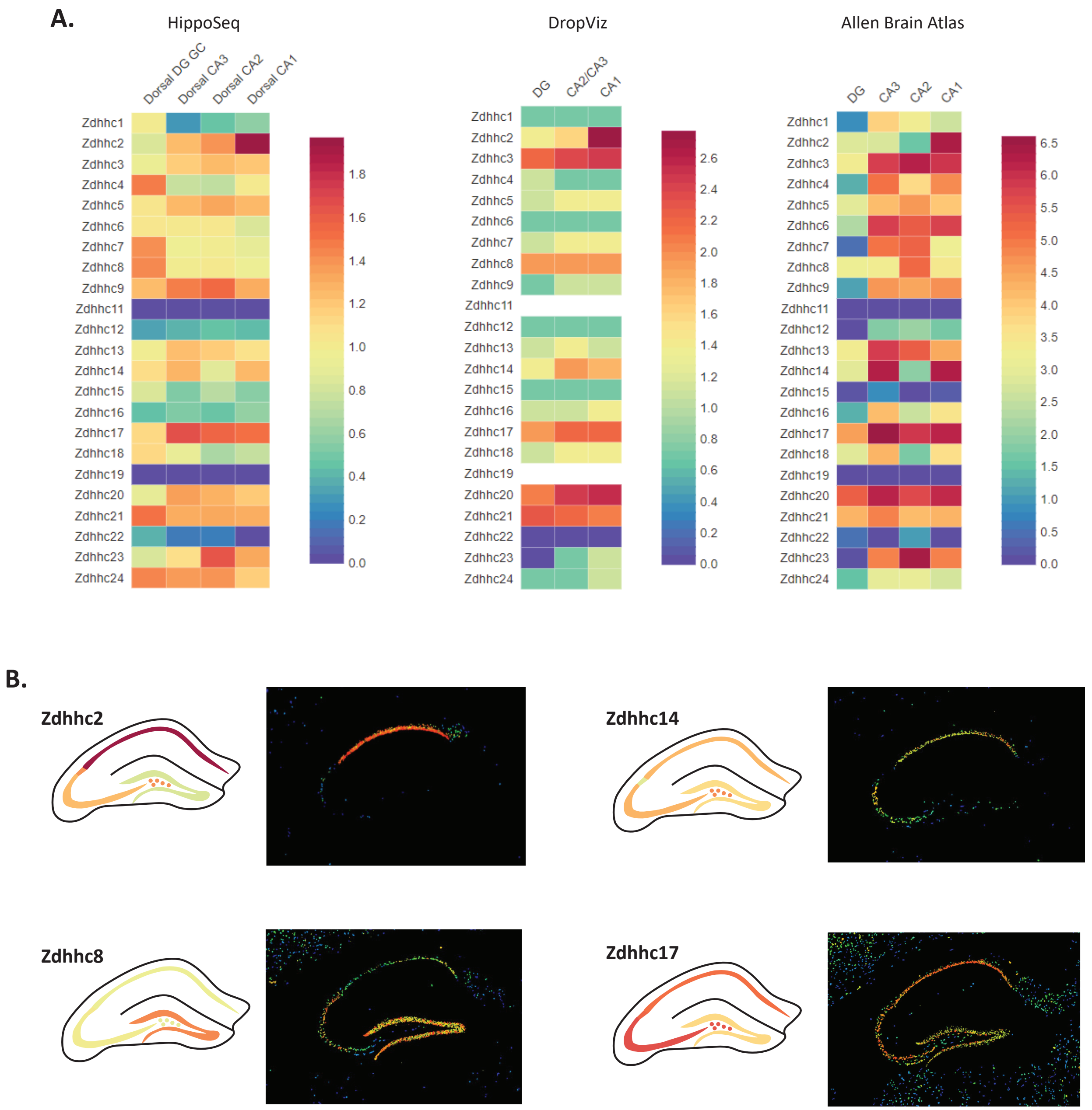


**Figure 2-figure supplement 1. Heterogeneous ZDHHC expression in excitatory neurons of the hippocampus**

(A) Heatmaps showing ZDHHC expression data from BrainPalmSeq for excitatory neurons of indicated subregions of the hippocampus. Original data are from 'Hipposeq' (Cembrowski et al., 2016; units: FPKM), 'DropViz' (Saunders et al., 2018; units: mean log2(counts per 10,000 + 1)) and Allen Mouse Brain 10X atlas (Yao et al., 2021; units: Trimmed Mean (25%-75%) Log2(CPM+1)).

(B) Expression maps from Figure 2A with in-situ hybridization images from Allen Brain ISH.

Figure 3-figure supplement 1

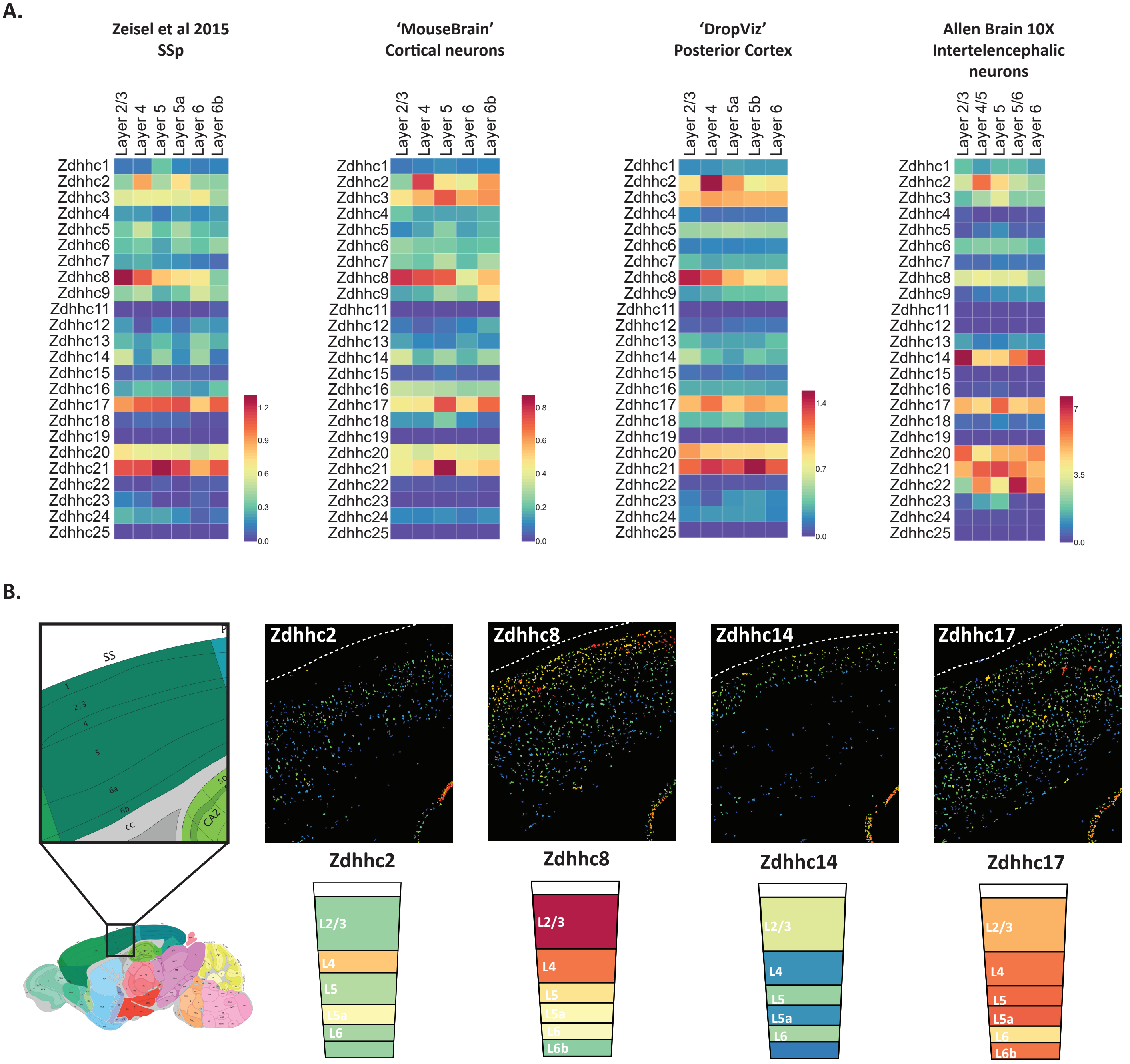


**Figure 3-figure supplement 1. Heterogeneous ZDHHC expression in excitatory neurons of the cortex**

(A) Heatmaps showing ZDHHC expression data from BrainPalmSeq for excitatory neurons of indicated layers of the cortex. Original data are from somatosensory cortex (Zeisel et al., 2015; units: mean log2(counts per 10,000 + 1)); 'MouseBrain' (Zeisel et al., 2018; mean log2(counts per 10,000 + 1); 'DropViz' posterior cortex (Saunders et al., 2018; units: mean log2(counts per 10,000 + 1)) and Allen Mouse Brain 10X atlas intertelencephalic neurons (Yao et al., 2021; units:  Trimmed Mean (25%-75%) Log2(CPM+1)).

(B) Expression maps from Figure 3A with in-situ hybridization images of somatosensory cortex from Allen Brain ISH.

Figure 4-figure supplement 1

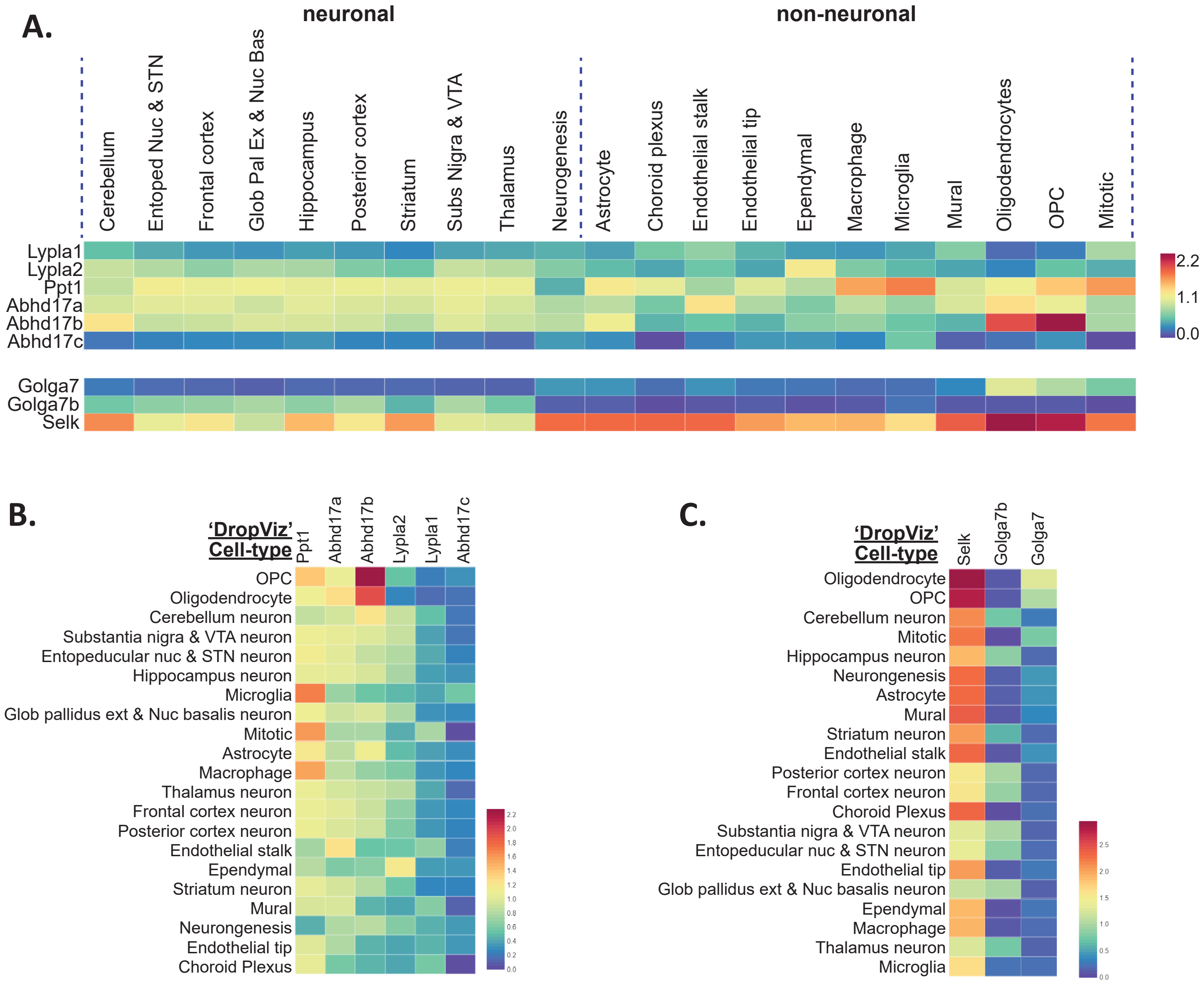


**Figure 4-figure supplement 1. Heterogeneous de-palmitoylating enzyme and accessory protein expression in mouse brain**

(A) Heatmap showing expression for the 6 depalmitoylating enzyme genes and 3 accessory protein genes extracted from 'DropViz' scRNAseq study of mouse brain (Saunders et al., 2018). Each column represents metacell averages. Metacells are organized along x-axis according to brain region (neuronal) or class (non-neuronal). Full metadata for this study available on BrainPalmSeq.

(B) Heatmap showing mean expression of the 6 depalmitoylating enzyme genes per brain region/class, with columns and rows sorted by descending mean gene expression per row/column.

(C) Heatmap showing mean expression of the 3 accessory protein genes per brain region/class, with columns and rows sorted by descending mean gene expression per row/column.

Figure 5-figure supplement 1

**A.**

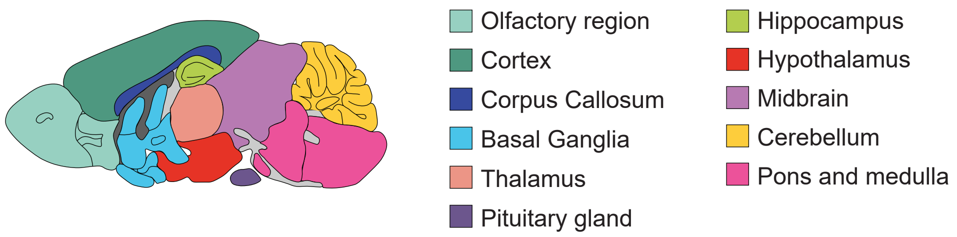


**Figure 5-figure supplement 1. Anatomical sampling for bulk RNAseq study**

(A) Diagram illustrating anatomical brain regions for bulk RNA expression data from Figure 5 (Sjöstedt et al., 2020).

Figure 6-figure supplement 1

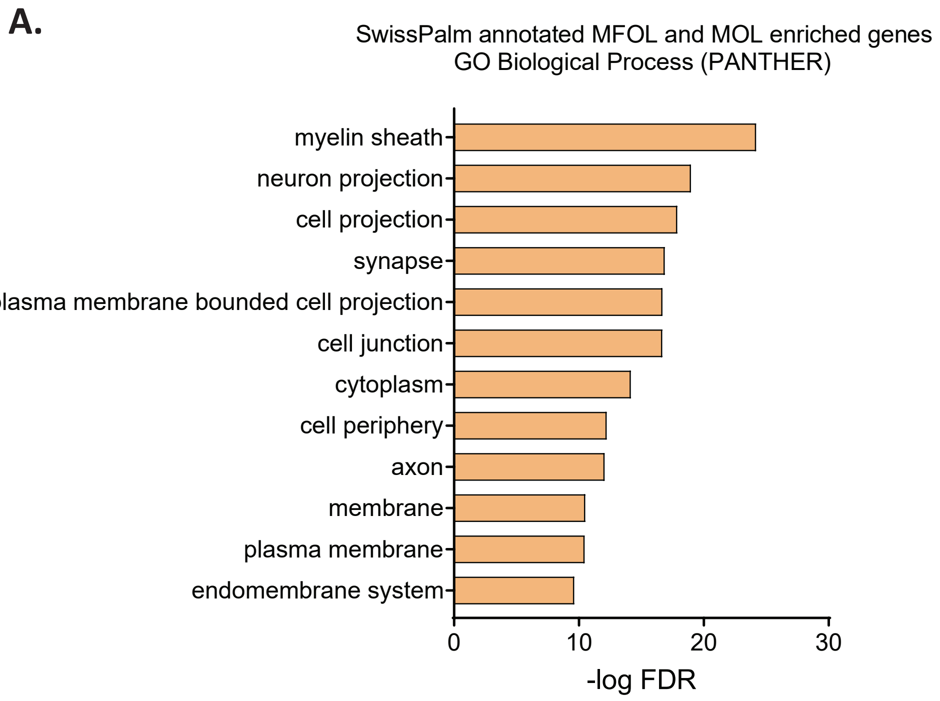


**Figure 6-figure supplement 1. GO cellular component analysis for oligodendrocyte enriched genes**

(A) Graph of GO cellular component analysis. Gene IDs from the full Marques oligodendrocyte dataset (Marques et al., 2016) that were enriched in MFOL and MOL subtypes and were also Uniprot reviewed and SwissPalm annotated were used as input.
